# A novel subtype of pineal projection neurons expressing melanopsin share a common developmental program with classical projection neurons

**DOI:** 10.1101/712091

**Authors:** Dora Sapède, Clair Chaigne, Patrick Blader, Elise Cau

## Abstract

The zebrafish pineal organ is a photoreceptive structure containing two main neuronal populations (photoreceptors and projections neurons). Here we describe a new pineal cell type that harbors both characteristics of projection neurons and photoreceptors. Indeed, a subpopulation of projection neurons expresses the melanopsin gene *opn4xa* suggesting photoreceptive properties. This population of hybrid cell fates, share a similar behaviour regarding dependency for BMP and Notch signalling pathways with classical non-photosensitive projection neurons (PNs) suggesting they are closer to the PN population. We describe two distinct types of activity within PNs: an achromatic LIGHT OFF activity in *opn4xa*^−^ PNs and a LIGHT ON activity elicited by green and blue light in *opn4xa*+ PNs. Altogether the discovery and characterization of *opn4xa*+ PNs suggest a previously unanticipated heterogeneity in the projection neuron population.

## Introduction

The zebrafish pineal gland is a neuroendocrine structure of the dorsal diencephalon which plays a key role in mediating the effects of the circadian clock on sleep/wake rhythms. An important part of its activity is due to pineal photoreceptors (PhRs) secreting the sleep-promoting hormone melatonin (Ben-Moshe Livne et al., 2016; Gandhi et al., 2015). PhR is an heterogeneous cell population that contains three described subtypes: a population of *red cone opsin* expressing cells, a population of *exorhodopsin* expressing cells (Cau et al., 2019; Clanton et al., 2013) as well as a population of cells coexpressing *parietopsin* (a green-sensitive opsin) and the UV-sensitive opsin, *parapinopsin1* (Cau et al., 2019; Koyanagi et al., 2015; Wada et al., 2018). In addition to PhRs, the larval pineal gland contains projection neurons (PNs) which, from early stages, project to the ventral diencephalon (Wilson and Easter, 1991) as well as a population of AgRP2+ cells which do not express neuronal markers nor exhibit a neuronal morphology and could share molecular characteristics with retinal-pigment epithelium cells (Shainer et al., 2017, 2019). Electrophysiological experiments in lampreys and goldfishes suggest that PNs receive and integrate inputs from the PhRs (Meissl et al., 1986; Uchida et al., 1992). While a significant amount of knowledge has been gained concerning the subpopulations of pineal PhRs and their function, much remains to be discovered concerning a possible corresponding heterogeneity in the PN population and PN functions are presently unknown. Most projection neurons are thought to function in a LIGHT OFF fashion. Indeed, electrophysiological experiments performed in the rainbow trout suggest that most teleost pineal projection neurons fire constantly in the dark and exhibit a LIGHT OFF response. This response is elicited at all visible wavelengths although it shows maximum efficiency at specific wavelengths. In addition, it is not clear whether all projection neurons respond to all wavelengths. Nevertheless, this response was referred to as ‘achromatic’ meaning that it is elicited by a wide range of wavelengths (Dodt, 1963; Morita, 1966). In contrast, in the same species, few projection neurons show a chromatic response. This response is excitatory at medium and long wavelengths and inhibitory in the violet-UV range (Morita, 1966). Achromatic LIGHT OFF responses were also recorded in other species such as frog, pike and turtle (Falcón and Meissl, 1981; Meissl and Ueck, 1980; Morita and Dodt, 1965) while chromatic LIGHT ON responses were described in lizards, frogs and pikes (Dodt and Heerd, 1962; Dodt and Meissl, 1982; Meissl and Donley, 1980). These results suggest heterogeneity in the population of pineal projection neurons in a number of species; it remains however to be discovered whether a similar heterogeneity exists in zebrafish as well as the molecular mechanisms that allow different types of PN to function so differently.

Heterogeneity also exists within the projection neurons population of the retina, the so-called RGCs (for Retinal Ganglion Cells; see Sanes and Masland, 2015 for a review). In particular, a subpopulation of RGCS has been shown to be directly photosensitive, owing to the expression of melanopsin, a blue-green photosensitive pigment. These atypical photoreceptors, referred to as ipRGCs (for intrinsically photosensitive RGCs), integrate light information from rods and cones as well as from their innate photosensitivity (Lucas et al., 2012) to regulate photo-entrainment of the circadian system. IpRGCs thus function both as photoreceptors and projection neurons.

In this paper, we describe the expression of one of the melanopsin genes: *opn4xa*, in a subset of PNs. PhRs and PNs can be distinguished by specific molecular markers as well as their requirement for signalling pathways. Indeed, the BMP (Bone Morphogenetic Protein) pathway is necessary and sufficient to trigger a PhR fate as well as to activate the Notch pathway in these cells; this latter being involved in inhibiting a PN fate (Cau and Blader, 2009; Cau et al., 2008; Quillien et al., 2011; Sapède and Cau, 2013). Analysis of embryos deficient for either the BMP or the Notch signalling pathways suggest that Notch is required to inhibit the production of *opn4xa*+ cells to the same extent than the rest of the PNs and that the BMP pathway is dispensable for its formation. These results suggest that despite exhibiting both characteristics of PNs and PhRs (the expression of an opsin), *opn4xa*+ cells are closer to PNs with which they share a developmental program. In addition, although the Wnt effector *tcf7* is specifically expressed in these cells, embryos exhibiting either a gain or a loss of Wnt activity show normal numbers of *opn4xa*+ cells. Finally, monitoring of the induction of the immediate early gene *c-fos* in different illumination regimes highlight two distinct activities in the PN population. A subpopulation of *opn4xa*- cells show a LIGHT OFF response when the embryos are adapted in white, blue, red or green light. In contrast, *opn4xa*+ PNs exhibit a LIGHT ON response following a pulse of white, blue or green light. Given the spectral sensitivity of their response to light, *opn4xa* + PNs could be directly photosensitive; we therefore draw a parallel between these cells and the ipRGCs of the retina.

## Results

### A subset of pineal projection neurons expresses the melanopsin gene *opn4xa*

In an effort to describe photoreceptor cell fates in the pineal, we looked at the expression of opsin photo-pigments. This approach led us to identify three classical photoreceptor cell types (Cau et al., 2019) as well as a new population of *opn4xa* + cells. Indeed, *opn4xa*, one of the five zebrafish melanopsin expressing genes (Matos-Cruz et al., 2011), was detected in a few cells located on both the left and right borders of the pineal organ (Fig. 1A). We used *in situ* hybridization combined with immunostaining against GFP in two transgenic lines Tg*(aanat2:GFP)*^*y8*^, Tg*(elavl3:EGFP)*^*knu3*^+ cells that labels PhR and PN respectively (Fig 1B, (Cau et al., 2008)) to unravel the nature of *opn4xa*+ cells. These experiments revealed that *opn4xa* expression is excluded from classical PhR (Fig. 1C,D), but present in PNs (Fig. 1E,F). *opn4xa* starts to be expressed in the pineal in 1-2 Tg*(elavl3:EGFP)*^*knu3*^ + cells at 22-23 hpf; at 48 hpf an average of 4 *opn4xa*+ cells is observed and this number is stable up to at least 10 dpf (Fig. 1I). To further characterize *opn4xa +* cells, we screened for different markers displaying restricted expression within the pineal organ. We observed that the gene encoding the Wnt transcriptional effector Tcf7 (T-cell specific transcription factor-7, (Veien et al., 2005) is expressed in a small subset of PNs (Fig S1 A-C). We took advantage of a *tcf7* enhancer trap line *Et(T2KHG)*^*nns8*^ (Nagayoshi et al., 2008) to further characterize *tcf7* expression in the pineal. We first confirmed that expression of GFP from the *Et(T2KHG)*^*nns8*^ enhancer trap co-localized with *tcf7* endogenous expression in PNs although this reporter line lacks expression in other epithalamic structures (Fig. S1 A-C). We next looked at co-labelling of the *Et(T2KHG)*^*nns8*^ enhancer trap line with *opn4xa* mRNA at various stages ranging from 24 hpf to 4 days. Up to stage 30 hpf included, the vast majority of GFP+ cells were also *opn4xa* + cells (Fig. 1 G,H, Fig S1 D-G). In contrast, from 2 days onwards, a population of cells showing a lower level of GFP appeared in the pineal of *Et(T2KHG)*^*nns8*^. Altogether, our results highlight the existence of a subpopulation of PNs expressing *opn4xa* that can specifically be labelled with the *Et(T2KHG)*^*nns8*^ enhancer trap line up to stage 30 hpf.

**Figure 1:**
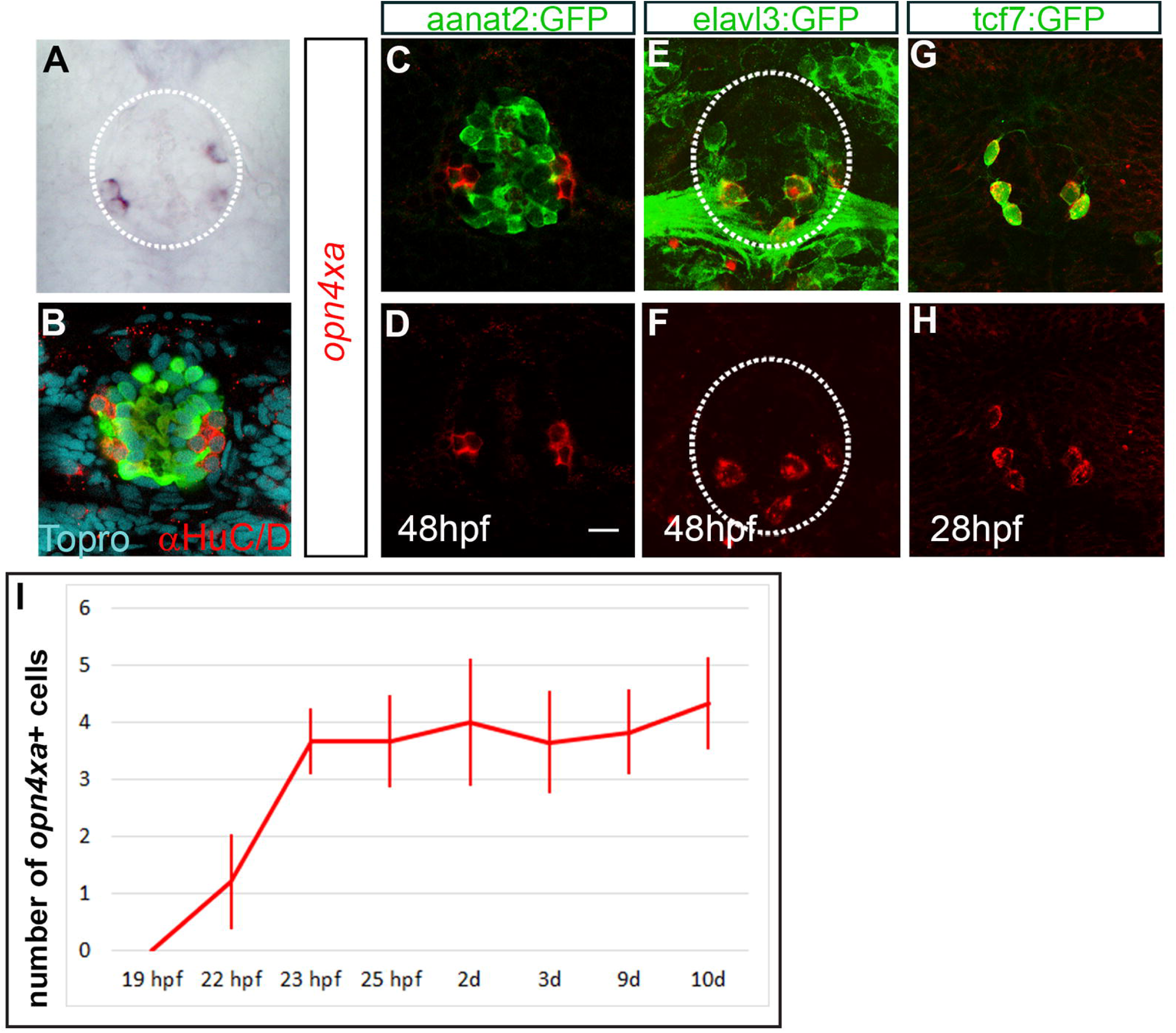
*opn4xa* is expressed in a restricted population of projection neurons within the pineal gland. (A) *in situ* hybridization at 48hpf showing pineal *opn4xa*+ cells. The pineal is highlighted with a white dotted circle. (B) Confocal projection showing the relative distribution of PNs (red, labelled with an anti-HuC/D antibody) and PhRs (green, labelled with an anti-GFP in a Tg*(aanat2:GFP)*^*y8*^ background) at 48 hpf. Topro (cyan) labels cell nuclei. (C-E) Co-expression of *opn4xa* mRNA (in red) on the one hand with the Tg*(aanat2:GFP)*^*y8*^ (‘aanat2:GFP’, C,D), the Tg*(elavl3:EGFP)*^*knu3*^ (‘elavl3:EGFP’, E,F) and the *Et(T2KHG)*^*nns8*^ (‘tcf7:GFP’, G,H transgenes (in green). (A-H) Anterior is upwards. Scale bars: 10 µm. (I)Time course of *opn4xa* expression in the developing pineal organ. Average numbers of *opn4xa*+ cells are quantified at different time points indicated in hours post-fertilization (hpf). Values are mean ±SD, n=4, 5, 3, 6, 9, 26, 6 and 6, respectively.

### Specification of the *opn4xa* + pineal fate does not require Wnt signalling

The fact that the *tcf7* enhancer trap line *Et(T2KHG)*^*nns8*^ is expressed in *opn4xa*+ cells prompted us to investigate whether Wnt signalling was involved in the specification of the *opn4xa*+ fate. First, we tested the expression of *opn4xa* mRNA in *Et(T2KHG)*^*nns8*^ homozygous embryos as this enhancer trap abolishes expression of the endogenous *tcf7* gene (Nagayoshi et al., 2008). Pineal *opn4xa+ cells* were observed in normal numbers in GFP+ embryos coming from *Et(T2KHG)*^*nns8*^ +/-incrosses suggesting that *tcf7* is not required for their specification (Fig 2 A,B). To look at a possible redundancy with other Wnt effectors, we analysed the expression of *tcf7l2* and *lef1* in the pineal and compared it with the *Et(T2KHG)*^*nns8*^ enhancer trap. Only the former appears expressed in the pineal (Figure S2) but do not show co-expression with the *tcf7* enhancer trap. Finally, we modulated Wnt activity at the time of the PN last division, using conditional overexpression of Dkk1 or Wnt8 in heat-shocked Tg*(hsp70l:dkk1b-GFP)*^w32^ and Tg*(hsp70:wnt8-GFP)*^*w34*^ embryos respectively (Stoick-Cooper et al., 2007). Heat-shocked transgenic embryos for the Tg*(hsp70l:dkk1b-GFP)*^w32^ transgene show normal expression of *opn4xa* (Fig 2B) suggesting that Wnt activity is dispensable for the specification of *opn4xa*+ cells. On the other hand, heat–shocked embryos transgenic for the Tg*(hsp70:wnt8-GFP)*^*w34*^ transgene show normal numbers of *opn4xa* + cells suggesting that Wnt activity is not either sufficient to drive the *opn4xa*+ fate. Therefore both loss and gain of function experiments suggest that specification of the *opn4xa*+ fate do not require Wnt signaling, as judged by the analysis of *opn4xa* expression.

**Figure 2:**
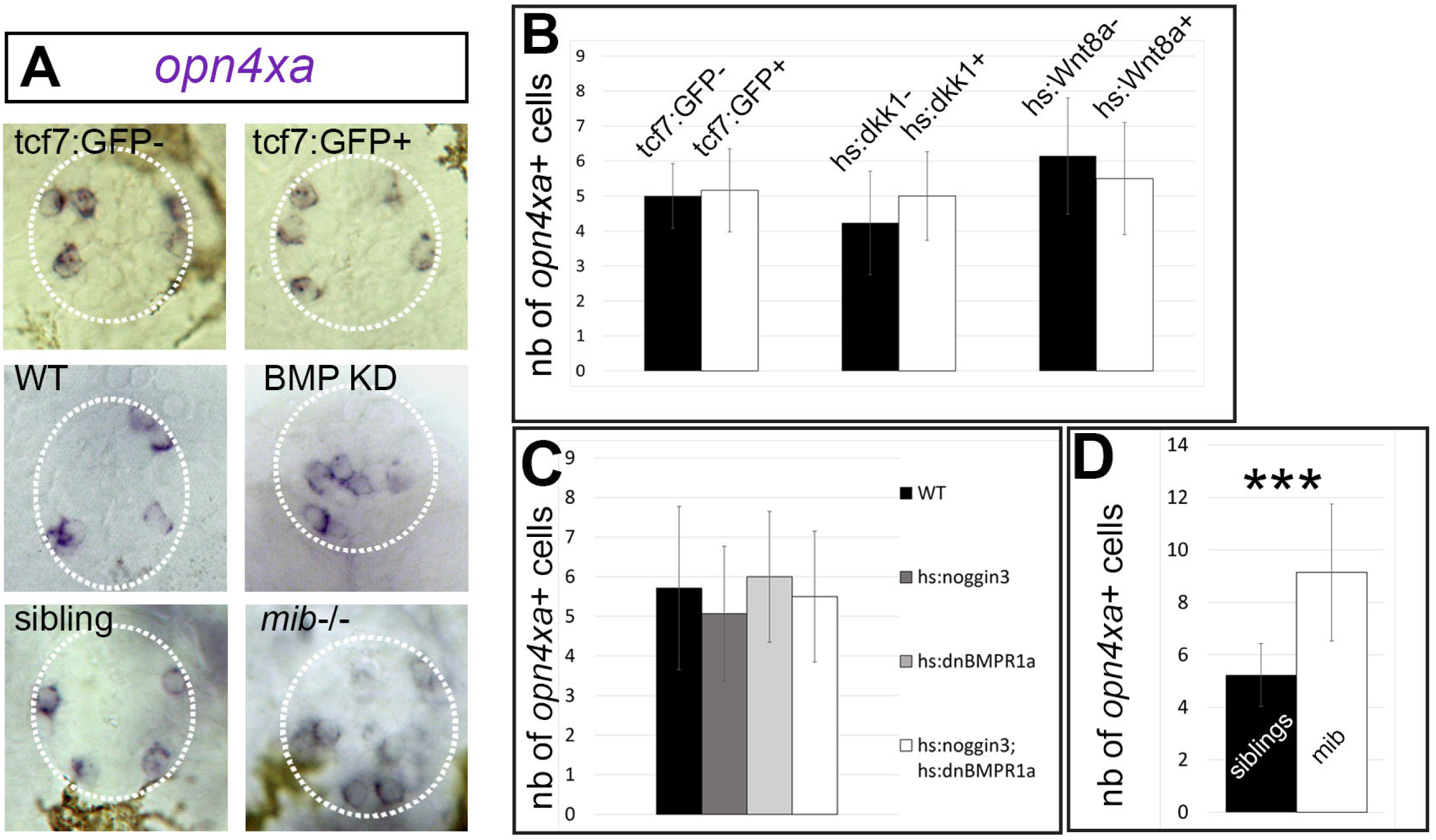
Effect of manipulating Notch, BMP and Wnt pathways on the *opn4xa+* fate. (A) Representative pictures of *in situ* hybridizations for *opn4xa* in the pineal organ upon BMP, Notch and Wnt modulation (as indicated) and their corresponding controls. (B-D) Average numbers of pineal *opn4xa+* positive cells upon manipulation of Wnt (B), BMP (C) or Notch signalling activity (D). Values indicated on the graphs are mean ±SD (error bars). Two tailed Mann Whitney non parametric tests, were used for B and D. *P<0.05; ***P<0.001. (B) Effect of gain or loss of Wnt activity on the number of pineal *opn4xa+* cells: Embryos from *Et(T2KHG)*^*nns8*^ +/-incrosses were assessed for GFP fluorescence; countings were performed at at 30hpf. The size of the GFP- and GFP+ populations (‘*tcf7:GFP-*’ and ‘*tcf7:GFP+*’) were respectively n=31 and n=8. Embryos carrying the Tg*(hsp70l:dkk1b-GFP)*^w32^ transgene after an heat shock at 18 hpf, were compared with negative siblings at 48 hpf (n=13 for the ‘*hs:dkk1*-’ and n=11 for the ‘*hs:dkk1*+’). Embryos carrying the Tg*(hsp70:wnt8-GFP)*^*w34*^ transgene (n=14) after an heat shock at 14 hpf, were compared to siblings (n=13) at 48 hpf. (C) Effect of a reduction in Notch activity: Control embryos (n=30) were compared to homozygous *mindbomb* (*mib-/-; n=7*) mutants at 48 hpf (D) Effect of a reduced BMP signalling: Embryos transgenic for Tg*(hsp70l:dnXla.Bmpr1a-GFP)*^*w30*^ (*hs:dnBMPR1a, n=12*) or Tg*(hsp70l:nog3)*^*fr14*^ (*hs:noggin3*, n=30) or both (n=18) were compared to siblings (n=7). Kruskal-Wallis test with Dunn’s multiple comparisons test, P=0.44; not significant.

### Specification of *opn4xa*+ cells do not require BMP signalling and is inhibited by Notch signalling

Classical PhRs are specified by an interplay of Notch and BMP activities with BMP activity being necessary and sufficient to activate the PhR fate and Notch being required within these cells to inhibit the inappropriate PN program (Cau and Blader, 2009; Cau et al., 2008; Quillien et al., 2011). Since *opn4xa*+ cells represent a mixed identity (hybrid) fate with both characteristics of a PhR and of a PN, we analyzed the effects of reducing BMP or Notch activities on the specification of these cells.

We used Tg*(hsp70l:dnXla.Bmpr1a-GFP)*^*w30*^; Tg*(hsp70l:nog3)*^*fr14*^ double transgenics heat-shocked at 14hpf to reduce BMP signalling within the pineal organ; indeed, such a condition was shown to reduce the number of Tg*(aanat2:GFP)*^*y8+*^ PhRs (Quillien et al., 2011). In contrast, the number of *opn4xa+* cells was similar in such Tg*(hsp70l:dnXla.Bmpr1a-GFP)*^*w30*^; Tg*(hsp70l:nog3)*^*fr14*^ double transgenic heat-shocked embryos compared to wildtype embryos (Fig 2A, C).

The effects of knocking down Notch activity was analyzed in the *mindbomb* (*mib*) mutant; as the *mib* gene, which encodes for a ring ubiquitin ligase, is required for activation of the Notch signalling pathway (Itoh et al., 2003). We noticed that *mib* embryos exhibit twice the amount of *opn4xa*+ cells at 48 hpf (Fig 2A,D). As our previous studies show that the total number of PN increases in the same proportion in *mib-/-* compared to wildtype embryos (Cau et al., 2008), we concluded that Notch exerts a similar inhibition on the *opn4xa*+ and the *opn4xa*-PN fate. Altogether with the results obtained in BMP knock down embryos, it suggests that pineal *opn4xa+* cells share with other pineal PNs, a similar absence of requirement for BMP activity for their specification, as well as a similar inhibition by Notch activity.

### The zebrafish pineal gland contains LIGHT ON and LIGHT OFF projection neurons and *opn4xa* + projection neurons respond to light in a LIGHT ON fashion

Electrophysiological experiments performed in the rainbow trout, frog, pike and turtle have shown that projection neurons are continuously activated in the dark (Dodt, 1963; Falcón and Meissl, 1981; Meissl and Ueck, 1980; Morita, 1966; Morita and Dodt, 1965) suggesting that they function in a LIGHT OFF mode. To assess the response of pineal projection neurons in general and those expressing *opn4xa* in particular, we looked at the expression of *c-fos*, an immediate early gene that is extensively used as a neuronal activation read out (reviewed in Guzowski et al., 2005), after a pulse of 30 minutes of dark in light adapted embryos (Fig 3A). While light-adapted embryos show no *c-fos* expression in the pineal (Fig 3Bi), a pulse of dark induced expression of *c-fos* expression in 8.36 ± 2.13 laterally-located cells per embryo (from n=14 embryos) (Fig 3Bii). The same experiment performed in *Et(T2KHG)*^*nns8*^ as well as Tg*(elavl3:EGFP)*^*knu3*^ embryos showed that these LIGHT OFF cells are *elavl3:EGFP* +/*tcf7:gfp*- projection neurons and thus correspond to *opn4xa*-projection neurons (Fig 3C,D).

**Figure 3:**
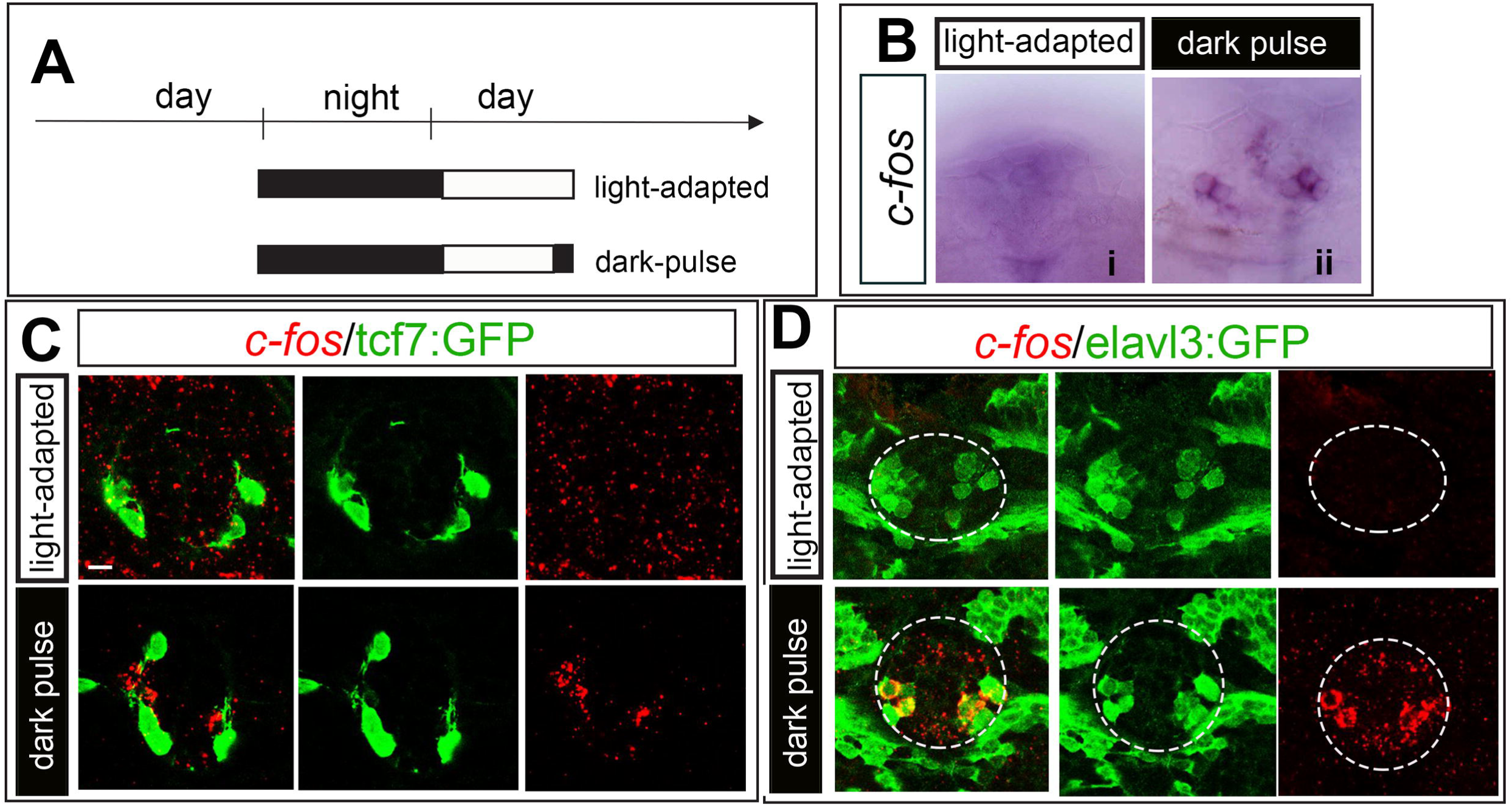
Identification of a population of *opn4xa*- PNs functioning in a LIGHT OFF mode: (A) Experimental paradigm for the ‘dark pulse versus light-adapted’ comparison Embryos were light-adapted for three hours before being submitted to a 30 min dark pulse (‘dark pulse’) or maintained in the light (‘light-adapted’) (B) Representative pictures of *c-fos* induction in the pineal organ after application of the light regimes described in A). i) light-adapted, ii) dark-pulse (C-E) Induction of *c-fos* in the different light regimes in embryos carrying the *Et(T2KHG)*^*nns8*^ (‘tcf7:GFP’, C) or the Tg*(elavl3:EGFP)*^*knu3*^ (‘elavl3:EGFP’, D) transgenes (in green). Scale bars: 10 µm. All embryos shown are 48 hpf.

On the other hand, a 30 min pulse of white light delivered to 2 day-old (2 dpf) dark adapted larvae induced *c-fos* expression in 3 ± 1,41 cells (from n=17 embryos) located laterally within the pineal organ (Fig 4Bii), whereas *c-fos* transcript was not detected in the pineal of non-light stimulated embryos (‘dark-adapted’, Fig 4Bi). Moreover, these cells strongly express the *Et(T2KHG)*^*nns8*^ transgene (Fig 4C). Similar results were obtained in 27 hpf and 3 dpf dark adapted embryos (Fig S3). These results suggest that a pulse of 30 min of white light specifically induce *c-fos* expression in *opn4xa* + projection neurons.

**Figure 4:**
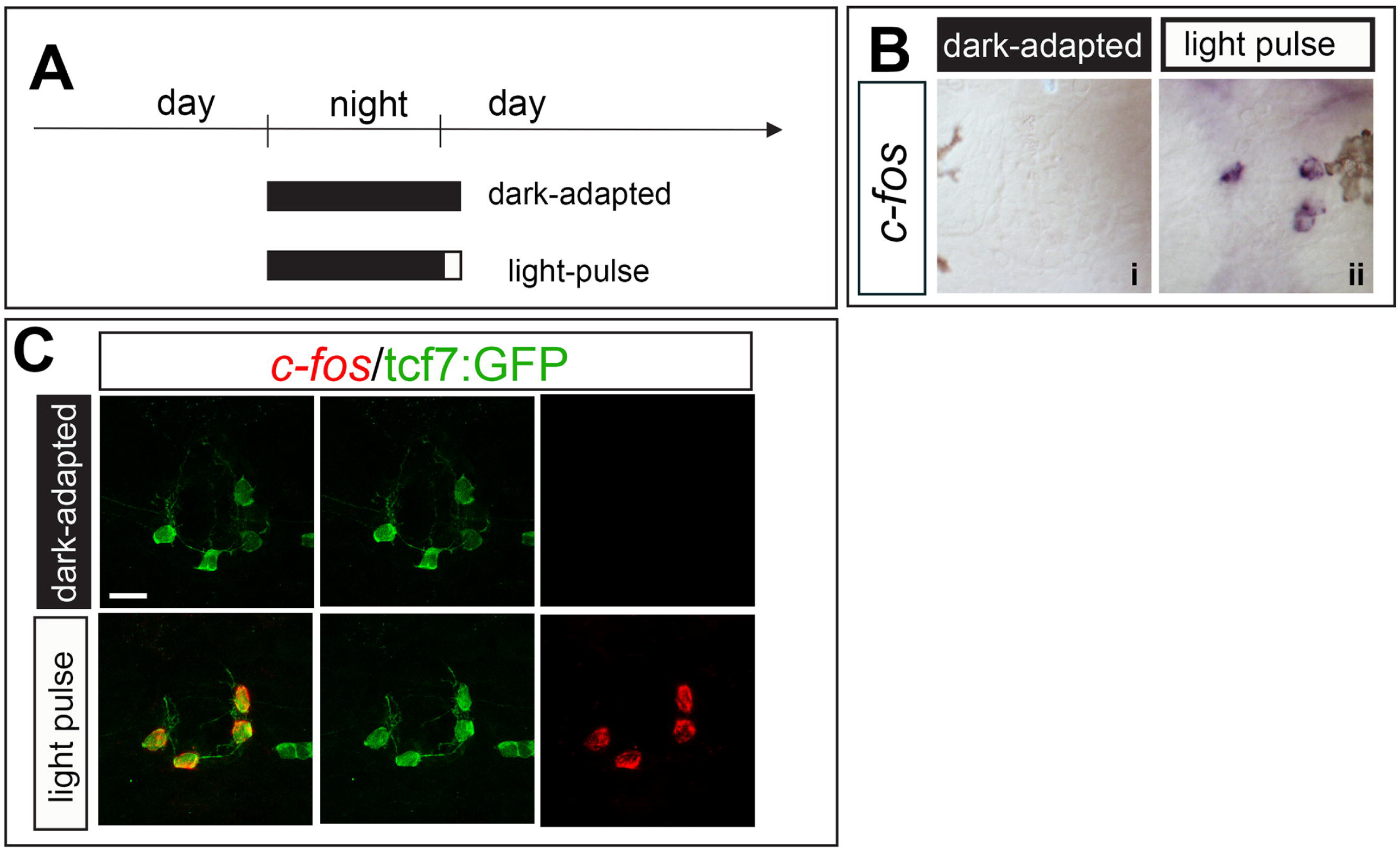
*opn4xa*+ PNs function in a LIGHT ON mode: (A) Experimental paradigm for the ‘light-pulse versus dark-adapted’ comparison Embryos were dark-adapted for more than 14 hours and submitted to a 30 min light pulse (‘light pulse’) or maintained in the dark (‘dark-adapted’) before fixation. (B) Representative pictures of *c-fos* induction in the pineal organ after application of the light regimes described in A). i) dark-adapted, ii) light-pulse (C-E) Induction of *c-fos* in the different light regimes in embryos carrying the *Et(T2KHG)*^*nns8*^ (‘tcf7:GFP’, C) transgene (in green). Scale bars: 10 µm.

Altogether, our results suggest an heterogeneity in the light-response within the PN population with *opn4xa*+ PNs functioning in a LIGHT ON fashion while *opn4xa*– PNs operate in a LIGHT OFF mode.

### LIGHT ON and LIGHT OFF responses are elicited by different range of wavelengths

Apart from OPN4XA, which is a blue-green light-sensitive photopigment (Davies et al., 2011), the pineal expresses opsin photo-pigments that are sensitive to green and red light (in classical photoreceptors, Cau et al., 2019). We next looked whether blue, red or green light were involved in the induction of *c-fos* expression. We observed that 30 mn light pulses in the blue and green range induce *c-fos* expression while red light could not induce a LIGHT ON response (Fig 5 A-F). Given that the absorbance spectrum of the OPN4XA photo-pigment overlaps with the emission of the green and blue LEDs used in this study; our interpretation is that this LIGHT ON response reflects the direct photosensitivity of *opn4xa*+ photo-pigment. In marked contrast with the results obtained for the LIGHT ON response, the LIGHT OFF response was elicited when the embryos have been adapted in either red, blue or green light (Figure 5G-L) suggesting that the LIGHT OFF response is elicited by a wide-range of wavelengths and can therefore be referred to as achromatic.

**Figure 5:**
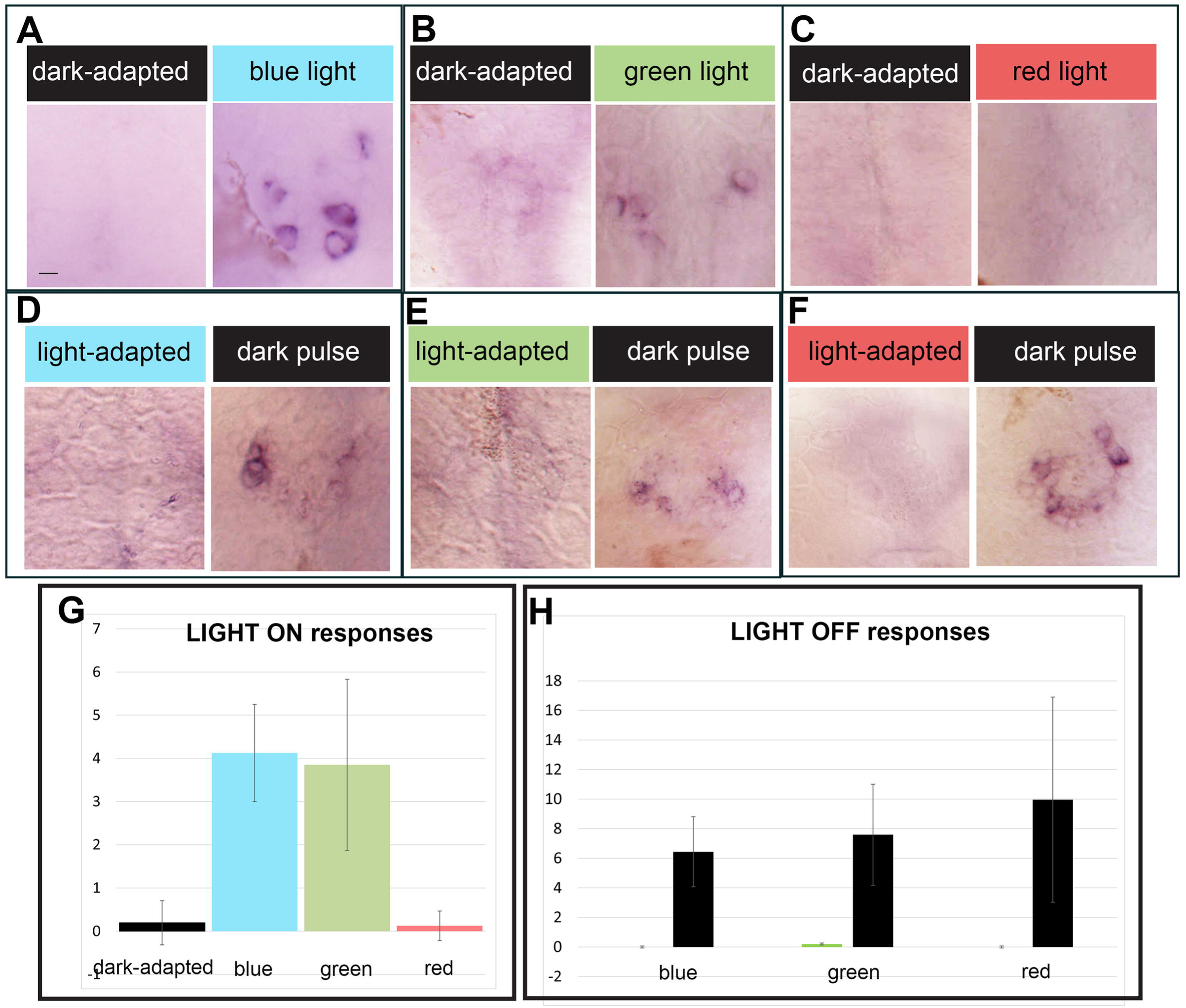
Different spectral sensitivities for the LIGHT ON and the LIGHT OFF responses within the pineal gland: (A-F) Representative pictures of *c-fos* induction in the pineal organ after application of light pulses in the blue, green or red range compared to dark adapted embryos (A-C) or after light adaptation with blue, green or red light followed by a dark pulse (‘dark pulse’) or maintained in the same lighting conditions (‘light-adapted’) (D-F) (G, H) Quantification of the number of cells activating c-fos after a light pulse (G, LIGHT ON responses) or a dark pulse (H, LIGHT OFF responses). G) Numbers of embryos : n= 41, 8, 20,16 respectively H) Numbers of embryos n=10,16,5,25,21,22 respectively Scale bar: 10 µm.

Altogether these results suggest that *opn4xa*– projection neurons function in a LIGHT OFF mode and most likely transduce the light input received by classical PhRs. In contrast, *opn4xa*+ cells are activated by light and most likely directly photosensitive.

## Discussion

In this paper, we describe a new population of *opn4xa*+ cells located in the zebrafish pineal gland. These cells express the green-blue photosensitive pigment OPN4XA and share several developmental characteristics with non-photosensitive projection neurons. In addition, we unveil two different light responses corresponding to two projection neurons’ populations:

-a LIGHT OFF response observed when light-adapted animals are submitted to a 30 min dark pulse. This response does not show chromatic specificity as it is elicited upon adaptation with either blue, green or red light and occurs in *opn4xa*– projection neurons.

-a LIGHT ON response observed in *opn4xa* + projection neurons upon illumination with blue or green but not red light

Altogether, these results suggest a previously unanticipated cellular diversity in the projection neuron population within the pineal gland.

### Decoupling of BMP and Notch activation does not contribute to the specification of *opn4xa* + fate

Projection neurons and photoreceptors of the pineal gland differ in their requirement for Notch and BMP signaling pathways: photoreceptors require BMP activation (while projection neurons do not) and BMP is in turn needed to activate the Notch pathway in these cells, which inhibits the projection neurons fate (Cau et al., 2008; Quillien et al., 2011). This led us to postulate that decoupling of the activation of BMP and Notch activity could represent an attractive mechanism to further diversify cell fates. Indeed, activation of the BMP signaling pathway without activation of Notch should lead to the generation of cells with a mixed photoreceptor/projection neuron identity (Cau and Blader, 2009). In contrast with this hypothesis, *opn4xa* + cells which exhibits both characteristics of photoreceptors and projection neurons, do not require BMP activity suggesting that they are more closely related to projection neurons than to classical photoreceptors. Along this line, Notch activity inhibits the production of *opn4xa* – and *opn4xa*+ projection neurons to the same extent (Fig 2) and therefore, neither BMP nor Notch activities contribute to specifying *opn4xa*+ versus *opn4xa* – PN fates.

### Wnt activity is dispensable for the specification of *opn4xa*+ PNs

Since *opn4xa* + and classical projection neurons show similar behavior upon BMP or Notch loss of function, what are the signals that could specify the *opn4xa*+ versus *opn4xa*- PN identity? The Wnt effector *tcf7* is specifically expressed in *opn4xa*+ PN and so is the *Et(T2KHG)*^*nns8*^ enhancer trap line at early stages (Figure S1, Figure 1). Nevertheless, *tcf7* mutants show a normal number of *opn4xa*+ pineal cells. Moreover, neither gain nor loss of global Wnt activity affects the *opn4xa* + fate (Fig 2), which suggests that Wnt activity is neither necessary nor sufficient for specification of *opn4xa* expression. It remains to be addressed whether *tcf7* (and/or Wnt signaling) is required to specify other aspects of the phenotype of these atypical PNs. Despite the absence of an observable function for *tcf7*, the *Et(T2KHG)*^*nns8*^ enhancer trap line provides a useful marker for *opn4xa* + cells at early stages which will help us analysing the projection pattern of these cells.

### *opn4xa* + PNs are the only described pineal projection neurons to function in a LIGHT ON mode in zebrafish

Our analysis confirms previous results obtained in another teleost the rainbow trout, as well as in numerous other species, showing that an important part of the projection neuron population function in a LIGHT OFF manner (Dodt, 1963; Falcón and Meissl, 1981; Meissl and Ueck, 1980; Morita, 1966; Morita and Dodt, 1965). Within the projection neuron population only *opn4xa*+ PNs function in a light ON fashion in response to visible light. Given that this light ON activity is only seen using blue and green light and not red light, it likely requires direct sensitivity from OPN4XA, a photo-pigment with an absorbance peak at 470 nm (Davies et al., 2011). This last result contrast with the observation that a LIGHT ON (excitatory response) is observed at medium to long wavelengths in the rainbow trout (Morita, 1966). A surprising degree of variation is indeed observed in the wavelengths triggering ON and OFF chromatic responses in different species. For instance, green light triggers the excitatory response in lizards and frogs while red light elicits this response in the pike. Similarly, the chromatic inhibitory response is stronger in the UV in fishes and frogs but mainly due to blue light in lizards (see Dodt and Meissl, (1982) for a review). It would be interesting to understand whether the LIGHT ON chromatic responses observed in lizard, frog and pike require a direct photosensitivity from specific projection neurons. Alternatively, in these species, pineal photoreceptors could elicit a dual excitatory/inhibitory action on their targets similar to what has been described for retinal photoreceptors which trigger a different response in bipolar cells depending on the type of glutamate receptor they express (Euler et al., 1996; Masu et al., 1995; de la Villa et al., 1995).

Are there other LIGHT ON projection neurons in the zebrafish pineal gland? Adaptation with white light allows for the identification of an average 8 LIGHT OFF PNs for around 3 LIGHT ON PNs. Since at 48 hpf the pineal gland contains 20 PNs, either some of the PN responses have not been identified or the LIGHT OFF cells do not systematically activate *c-fos* following a 30 min dark pulse, in the present testing conditions. It thus remains possible that there are other unidentified projection neurons that respond in a LIGHT ON fashion for instance in response to UV light.

### Cell-cell communication within the pineal and with its targets

While a complete description of the neurotransmitter/neuromodulator content of all zebrafish pineal neurons is still lacking, glycinergic cells with a photoreceptor morphology have been observed in the pineal gland (Marquart et al., 2015; Moly et al., 2014)). These observations support that at least a proportion of pineal photoreceptors could exert an inhibitory action on projection neurons, therefore providing a possible mechanism for the LIGHT OFF response we describe. Alternatively, photoreceptors could constantly produce an excitatory neurotransmitter in the dark, a process interrupted by light, which would lead to an interruption of projection neuron activity in the presence of light through a “disfacilitation mechanism” (Uchida et al., 1992). A more detailed analysis of the neurotransmitter/neuromodulator content of all pineal neurons and *opn4xa*+ PNs in particular would be useful. Finally, the fact that the LIGHT OFF response is observed independently of the wavelength at which the embryos are adapted suggests that LIGHT OFF projection neurons could receive inputs from several types of pineal photoreceptors.

### Towards an understanding for the role of *opn4xa* + PNs

Although a direct photosensitivity remains to be adressed, *opn4xa*+ PNs share many characteristics with retinal ipRGCS which raises intriguing questions regarding the possible functions of these pineal cells. Indeed, in mammals, ipRGCs exhibit crucial roles during the control of circadian, wake/sleep rhythms as well as in mediating the direct effects of illumination on physiology and behavior (see Lazzerini Ospri et al., (2017); Lucas et al.,(2012) for reviews). In particular, ipRGCs transmit light information, owing to its intrinsic photosensitivity as well as inputs from rods and cones, to the mammalian ‘master clock’ the suprachiasmatic nucleus (Hannibal and Fahrenkrug, 2004; Hattar et al., 2002, 2006). It is unclear whether a suprachiasmatic nucleus is present in the larval/embryonic zebrafish and whether *opn4xa*+retinal or pineal cells project onto such a structure. The zebrafish pineal gland, on the other hand has long been thought to play a ‘master clock’ role owing to its direct photosensitivity, its capacity to generate intrinsic rhythms of melatonin secretion as well as the fact that disrupting the function of the molecular clock in pineal PhRs affect circadian rhythms of locomotor activity (Ben-Moshe Livne et al., 2016; Bolliet et al., 1997; Cahill, 1996, 1997; Falcón et al., 1989). Could *opn4xa*+ PNs play a role alongside with pineal PhRs in the function of the circadian system? Analysis of an *opn4xa* mutant allele combined with retina-specific and/or pineal specific transgenic tools will help us understand whether *opn4xa*+ PNs are important players in this field.

## Experimental Procedures

### Zebrafish lines and developmental conditions

All animals were handled in the CBI fish facility, which is certified by the French Ministry of Agriculture (approval number A3155510). The project was approved by the French Ministry of Teaching and Research (agreement number APAFIS#3653-2016011512005922).

Embryos were reared at 28 degrees and staged according to standard protocols. All the transgenic and mutant lines have been described previously *Tg(aanat2:GFP)*^*y8*^ (Gothilf et al., 2002); Tg*(elavl3:EGFP)*^*knu3*^ (Park et al., 2000); *Et(T2KHG)*^*nns8*^ (Nagayoshi et al., 2008); Tg*(hsp70l:dkk1b-GFP)*^w32^ and Tg*(hsp70:wnt8-GFP)*^*w34*^ respectively (Stoick-Cooper et al., 2007); *Tg(hsp70l:dnXla.Rbpj-MYC)*^*vu21*^ and Tg*(hsp70l:nog3)*^*fr14*^ (Chocron et al., 2007); *mib*^*ta52b*^ (Itoh et al., 2003). Conditions of heat-shock were as follows: *Tg(hsp70l:dnXla.Rbpj-MYC)*^*vu21*^ and Tg*(hsp70l:nog3)*^*fr14*^ : 30 minutes at 39.5°C; Tg*(hsp70l:dkk1b-GFP)*^w32^ and Tg*(hsp70:wnt8-GFP)*^*w34* :^ 1 hour at 38°C.

#### Application of different light regimes

Illumination with white light was performed using a regular neon lamp (80 lux). Illumination with blue, green and red light was performed in a MWP unit (Zantiks) equipped with standard blue, green and red LEDs. The characteristics of the light emitted from these LEDS is as follows:

Blue light: 92 lux, λmax= 449nm, half-band width: 25,4nm

Green light: 109 lux, λmax= 512 nm, half-band width: 33,07 nm

Red light 34 lux, λmax= 627 nm, half-band width: 21,67 nm

Scripts encoding the different light regimes are available upon request.

### Immunohistochemistry and in situ hybridization

Immunohistochemistry and in situ hybridization were performed as previously described; in particular, fluorescent in situ hybridization was revealed using either Fast Red (Sigma) or the TSA Plus System (TSA-Fluorescein, Perkin Elmer; Cau et al., 2019). Immunohistochemistry against Tcf7l2 was performed as described in Hüsken et al., (2014) using an anti-Human TCF3,4 antibody (1:400, Biomol, clone 0.T.148).

Probes used in this study were as follows: *opn4xa* (Matos-Cruz et al., 2011), *c-fos* (Ellis et al., 2012), *lef1* and *tcf7* (Veien et al., 2005).

## Acknowledgements

We thank Serge Mazères for his help with the spectral analysis of LED light, Richard Dorsky for providing the *Et(T2KHG)*^*nns8*^; Tg*(hsp70l:dkk1b-GFP)*^w32^ and Tg*(hsp70:wnt8-GFP)*^*w34*^ transgenic lines as well as M. Halpern and K. Soanes for sharing their probes. We are indebted to members of the Blader team for helpful discussions. This work was supported by the Centre National de la Recherche Scientifique (CNRS); the Institut National de la Santé et de la Recherche Médicale (INSERM); Université de Toulouse III (UPS); Fondation pour la Recherche Médicale (FRM; DEQ20131029166); Fédération pour la Recherche sur le Cerveau (FRC); Association Rétina France and the Ministère de la Recherche. We would like to thank Brice Ronsin, Stéphanie Bosch and the Toulouse RIO Imaging platform; as well as Stéphane Relexans, Aurore Laire and Richard Brimicombe for taking care of the fish.

## Supplemental figure legends

**Supplemental Figure 1: Characterization of the expression driven by the *Et(T2KHG)***^***nns8***^ **(‘tcf7:GFP’) enhancer trap in the pineal gland :**

(A-F) Comparison of *tcf7* (A-C) or *opn4xa* (D-F) expression (in red, in situ hybridization) with GFP expression from the *Et(T2KHG)*^*nns8*^ (‘tcf7:GFP’) transgene (in green, immunohistochemistry). White arrowheads highlight co-expressing cells. White arrow show a weak tcf7:GFP+ *opn4xa* – cell. Scale bars: 10 µm. (G) Comparison of the number of cells expressing tcf7:GFP strongly or weakly and *opn4xa*. Means ± S.D are shown.

**Supplemental Figure 2: Comparison of the expression of *tcf7l2* and *lef1* with the *Et(T2KHG)***^***nns8***^ **(‘tcf7:GFP’) enhancer trap in the pineal gland**

(A-F) Expression of two other TCF encoding genes: *tcf7l2* (A-C) or *lef1* (D-F) (in red, in situ hybridization) with expression from the *Et(T2KHG)*^*nns8*^ (‘tcf7:GFP’) transgene (in green, immunohistochemistry). (G) Double immunostaining with anti-GFP (green) and anti-TCF7l2 (red) in pineal neurons.

Anterior is up. All embryos are 48 hpf. Scale bars: 10 µm.

**Supplemental Figure 3 : Induction of *c-fos* expression following a light pulse in the pineal gland at 27 hpf and 3 dpf.**

Expression of *c-fos* (in red, *in situ* hybridization) compared with expression from the *Et(T2KHG)*^*nns8*^ (‘tcf7:GFP’) transgene (in green, immunohistochemistry) at 27hpf (A-C) and 3dpf (D-F). Scale bars: 10 µm.

## References

Ben-Moshe Livne, Z., Alon, S., Vallone, D., Bayleyen, Y., Tovin, A., Shainer, I., Nisembaum, L.G., Aviram, I., Smadja-Storz, S., Fuentes, M., et al. (2016). Genetically Blocking the Zebrafish Pineal Clock Affects Circadian Behavior. PLoS Genet. 12, e1006445.

Bolliet, V., Bégay, V., Taragnat, C., Ravault, J.P., Collin, J.P., and Falcón, J. (1997). Photoreceptor cells of the pike pineal organ as cellular circadian oscillators. Eur. J. Neurosci. 9, 643–653.

Cahill, G.M. (1996). Circadian regulation of melatonin production in cultured zebrafish pineal and retina. Brain Res. 708, 177–181.

Cahill, G.M. (1997). Circadian melatonin rhythms in cultured zebrafish pineals are not affected by catecholamine receptor agonists. Gen. Comp. Endocrinol. 105, 270–275.

Cau, E., and Blader, P. (2009). Notch activity in the nervous system: to switch or not switch? Neural Develop. 4, 36.

Cau, E., Quillien, A., and Blader, P. (2008). Notch resolves mixed neural identities in the zebrafish epiphysis. Development. 135, 2391–2401.

Cau, E., Ronsin, B., Bessière, L., and Blader, P. (2019). A Notch-mediated, temporal asymmetry in BMP pathway activation promotes photoreceptor subtype diversification. PLoS Biol. 17, e2006250.

Chocron, S., Verhoeven, M.C., Rentzsch, F., Hammerschmidt, M., and Bakkers, J. (2007). Zebrafish Bmp4 regulates left-right asymmetry at two distinct developmental time points. Dev. Biol. 305, 577–588.

Clanton, J.A., Hope, K.D., and Gamse, J.T. (2013). Fgf signaling governs cell fate in the zebrafish pineal complex. Development. 140, 323–332.

Davies, W.I.L., Zheng, L., Hughes, S., Tamai, T.K., Turton, M., Halford, S., Foster, R.G., Whitmore, D., and Hankins, M.W. (2011). Functional diversity of melanopsins and their global expression in the teleost retina. Cell. Mol. Life Sci. CMLS 68, 4115–4132.

Dodt, E. (1963). PHOTOSENSITIVITY OF THE PINEAL ORGAN IN THE TELEOST, SALMO IRIDEUS (GIBBONS). Experientia 19, 642–643.

Dodt, E., and Heerd, E. (1962). Mode of action of pineal nerve fibers in frogs. J. Neurophysiol. 25, 405–429.

Dodt, E., and Meissl, H. (1982). The pineal and parietal organs of lower vertebrates. Experientia 38, 996–1000.

Ellis, L.D., Seibert, J., and Soanes, K.H. (2012). Distinct models of induced hyperactivity in zebrafish larvae. Brain Res. 1449, 46–59.

Euler, T., Schneider, H., and Wässle, H. (1996). Glutamate responses of bipolar cells in a slice preparation of the rat retina. J. Neurosci.16, 2934–2944.

Falcón, J., and Meissl, H. (1981). The Photosensory Function of the Pineal Organ of the Pike (Esox lucius L.) Correlation Between Structure and Function. J. Comp. Physiol. A 144, 127–137.

Falcón, J., Marmillon, J.B., Claustrat, B., and Collin, J.P. (1989). Regulation of melatonin secretion in a photoreceptive pineal organ: an in vitro study in the pike. J. Neurosci. 9, 1943–1950.

Gandhi, A.V., Mosser, E.A., Oikonomou, G., and Prober, D.A. (2015). Melatonin is required for the circadian regulation of sleep. Neuron 85, 1193–1199.

Gothilf, Y., Toyama, R., Coon, S.L., Du, S.-J., Dawid, I.B., and Klein, D.C. (2002). Pineal-specific expression of green fluorescent protein under the control of the serotonin-N-acetyltransferase gene regulatory regions in transgenic zebrafish. Dev. Dyn. 225, 241–249.

Guzowski, J.F., Timlin, J.A., Roysam, B., McNaughton, B.L., Worley, P.F., and Barnes, C.A. (2005). Mapping behaviorally relevant neural circuits with immediate-early gene expression. Curr. Opin. Neurobiol. 15, 599–606.

Hannibal, J., and Fahrenkrug, J. (2004). Melanopsin containing retinal ganglion cells are light responsive from birth. Neuroreport 15, 2317–2320.

Hattar, S., Liao, H.W., Takao, M., Berson, D.M., and Yau, K.W. (2002). Melanopsin-containing retinal ganglion cells: architecture, projections, and intrinsic photosensitivity. Science 295, 1065–1070.

Hattar, S., Kumar, M., Park, A., Tong, P., Tung, J., Yau, K.-W., and Berson, D.M. (2006). Central projections of melanopsin-expressing retinal ganglion cells in the mouse. J. Comp. Neurol. 497, 326–349.

Hüsken, U., Stickney, H.L., Gestri, G., Bianco, I.H., Faro, A., Young, R.M., Roussigne, M., Hawkins, T.A., Beretta, C.A., Brinkmann, I., et al. (2014). Tcf7l2 is required for left-right asymmetric differentiation of habenular neurons. Curr. Biol. CB 24, 2217–2227.

Itoh, M., Kim, C.-H., Palardy, G., Oda, T., Jiang, Y.-J., Maust, D., Yeo, S.-Y., Lorick, K., Wright, G.J., Ariza-McNaughton, L., et al. (2003). Mind bomb is a ubiquitin ligase that is essential for efficient activation of Notch signaling by Delta. Dev. Cell 4, 67–82.

Koyanagi, M., Wada, S., Kawano-Yamashita, E., Hara, Y., Kuraku, S., Kosaka, S., Kawakami, K., Tamotsu, S., Tsukamoto, H., Shichida, Y., et al. (2015). Diversification of non-visual photopigment parapinopsin in spectral sensitivity for diverse pineal functions. BMC Biol. 13, 73.

Lazzerini Ospri, L., Prusky, G., and Hattar, S. (2017). Mood, the Circadian System, and Melanopsin Retinal Ganglion Cells. Annu. Rev. Neurosci. 40, 539–556.

Lucas, R.J., Lall, G.S., Allen, A.E., and Brown, T.M. (2012). How rod, cone, and melanopsin photoreceptors come together to enlighten the mammalian circadian clock. Prog. Brain Res. 199, 1–18.

Marquart, G.D., Tabor, K.M., Brown, M., Strykowski, J.L., Varshney, G.K., LaFave, M.C., Mueller, T., Burgess, S.M., Higashijima, S.-I., and Burgess, H.A. (2015). A 3D Searchable Database of Transgenic Zebrafish Gal4 and Cre Lines for Functional Neuroanatomy Studies. Front. Neural Circuits 9, 78.

Masu, M., Iwakabe, H., Tagawa, Y., Miyoshi, T., Yamashita, M., Fukuda, Y., Sasaki, H., Hiroi, K., Nakamura, Y., and Shigemoto, R. (1995). Specific deficit of the ON response in visual transmission by targeted disruption of the mGluR6 gene. Cell 80, 757–765.

Matos-Cruz, V., Blasic, J., Nickle, B., Robinson, P.R., Hattar, S., and Halpern, M.E. (2011). Unexpected diversity and photoperiod dependence of the zebrafish melanopsin system. PloS One 6, e25111.

Meissl, H., and Donley, C.S. (1980). Change of threshold after light-adaptation of the chromatic response of the frog’s pineal organ (Stirnorgan). Vision Res. 20, 379–383.

Meissl, H., and Ueck, M. (1980). Extraocular Photoreception of the Pineal Gland of the Aquatic Turtle Pseudemys scripta elegans. J. Comp. Physiol. A 140, 173–179.

Meissl, H., Nakamura, T., and Thiele, G. (1986). Neural response mechanisms in the photoreceptive pineal organ of goldfish. Comp. Biochem. Physiol. A 84, 467–473.

Moly, P.K., Ikenaga, T., Kamihagi, C., Islam, A.F.M.T., and Hatta, K. (2014). Identification of initially appearing glycine-immunoreactive neurons in the embryonic zebrafish brain. Dev. Neurobiol. 74, 616–632.

Morita, Y. (1966). [Lead pattern of the pineal neuron of the rainbow trout (Salmo irideus) by illumination of the diencephalon]. Pflugers Arch. Gesamte Physiol. Menschen Tiere 289, 155–167.

Morita, Y., and Dodt, E. (1965). Nervous activity of the frog’s epiphysis cerebri in relation to illumination. Experientia 21, 221–222.

Nagayoshi, S., Hayashi, E., Abe, G., Osato, N., Asakawa, K., Urasaki, A., Horikawa, K., Ikeo, K., Takeda, H., and Kawakami, K. (2008). Insertional mutagenesis by the Tol2 transposon-mediated enhancer trap approach generated mutations in two developmental genes: tcf7 and synembryn-like. Development. 135, 159–169.

Park, H.C., Kim, C.H., Bae, Y.K., Yeo, S.Y., Kim, S.H., Hong, S.K., Shin, J., Yoo, K.W., Hibi, M., Hirano, T., et al. (2000). Analysis of upstream elements in the HuC promoter leads to the establishment of transgenic zebrafish with fluorescent neurons. Dev. Biol. 227, 279–293.

Quillien, A., Blanco-Sanchez, B., Halluin, C., Moore, J.C., Lawson, N.D., Blader, P., and Cau, E. (2011). BMP signaling orchestrates photoreceptor specification in the zebrafish pineal gland in collaboration with Notch. Development. 138, 2293–2302.

Sanes, J.R., and Masland, R.H. (2015). The types of retinal ganglion cells: current status and implications for neuronal classification. Annu. Rev. Neurosci. 38, 221–246.

Sapède, D., and Cau, E. (2013). The pineal gland from development to function. Curr. Top. Dev. Biol. 106, 171–215.

Shainer, I., Buchshtab, A., Hawkins, T.A., Wilson, S.W., Cone, R.D., and Gothilf, Y. (2017). Novel hypophysiotropic AgRP2 neurons and pineal cells revealed by BAC transgenesis in zebrafish. Sci. Rep. 7, 44777.

Shainer, I., Michel, M., Marquart, G.D., Bhandiwad, A.A., Zmora, N., Ben-Moshe Livne, Z., Zohar, Y., Hazak, A., Mazon, Y., Förster, D., et al. (2019). Agouti-Related Protein 2 Is a New Player in the Teleost Stress Response System. Curr. Biol. CB 29, 2009–2019.e7.

Stoick-Cooper, C.L., Weidinger, G., Riehle, K.J., Hubbert, C., Major, M.B., Fausto, N., and Moon, R.T. (2007). Distinct Wnt signaling pathways have opposing roles in appendage regeneration. Development. 134, 479–489.

Uchida, K., Nakamura, T., and Morita, Y. (1992). Signal transmission from pineal photoreceptors to luminosity-type ganglion cells in the lamprey, Lampetra japonica. Neuroscience 47, 241–247.

Veien, E.S., Grierson, M.J., Saund, R.S., and Dorsky, R.I. (2005). Expression pattern of zebrafish tcf7 suggests unexplored domains of Wnt/beta-catenin activity. Dev. Dyn. Off. Publ. Am. Assoc. Anat. 233, 233–239.

de la Villa, P., Kurahashi, T., and Kaneko, A. (1995). L-glutamate-induced responses and cGMP-activated channels in three subtypes of retinal bipolar cells dissociated from the cat. J. Neurosci. 15, 3571–3582.

Wada, S., Shen, B., Kawano-Yamashita, E., Nagata, T., Hibi, M., Tamotsu, S., Koyanagi, M., and Terakita, A. (2018). Color opponency with a single kind of bistable opsin in the zebrafish pineal organ. Proc. Natl. Acad. Sci. U. S. A. 115, 11310–11315.

Wilson, S.W., and Easter, S.S. (1991). Stereotyped pathway selection by growth cones of early epiphysial neurons in the embryonic zebrafish. Development. 112, 723–746.

