## Supplementary figures and images for "A novel subtype of pineal projection neurons expressing melanopsin share a common developmental program with classical projection neurons"

### Supplemental Figure 1

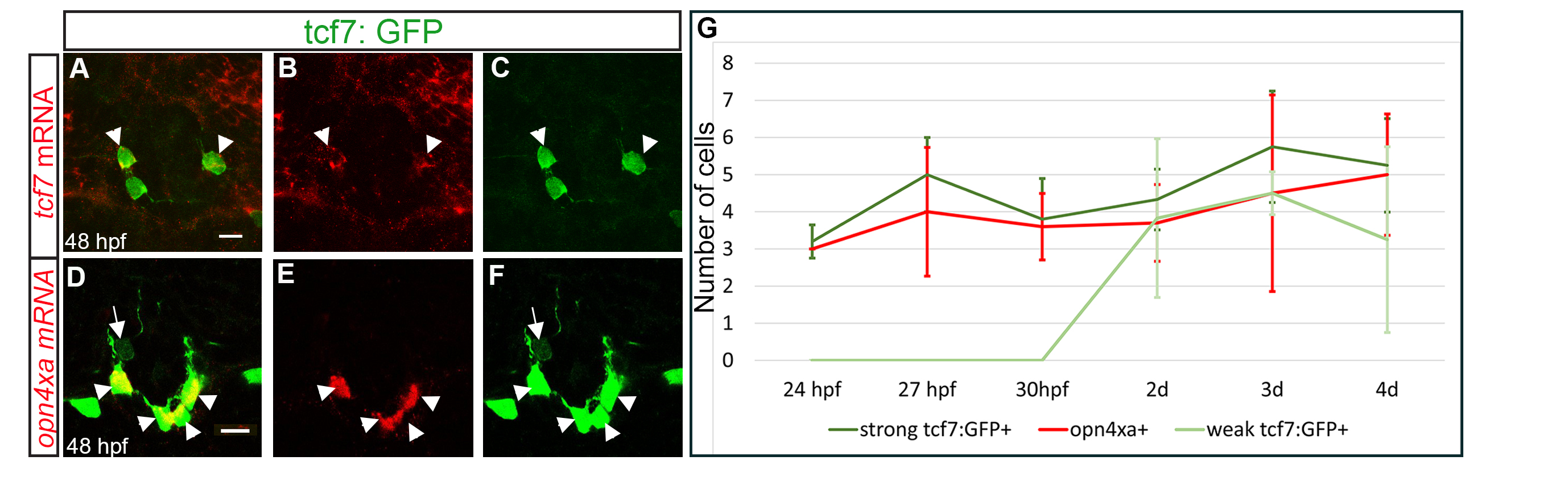

### Supplemental Figure 2

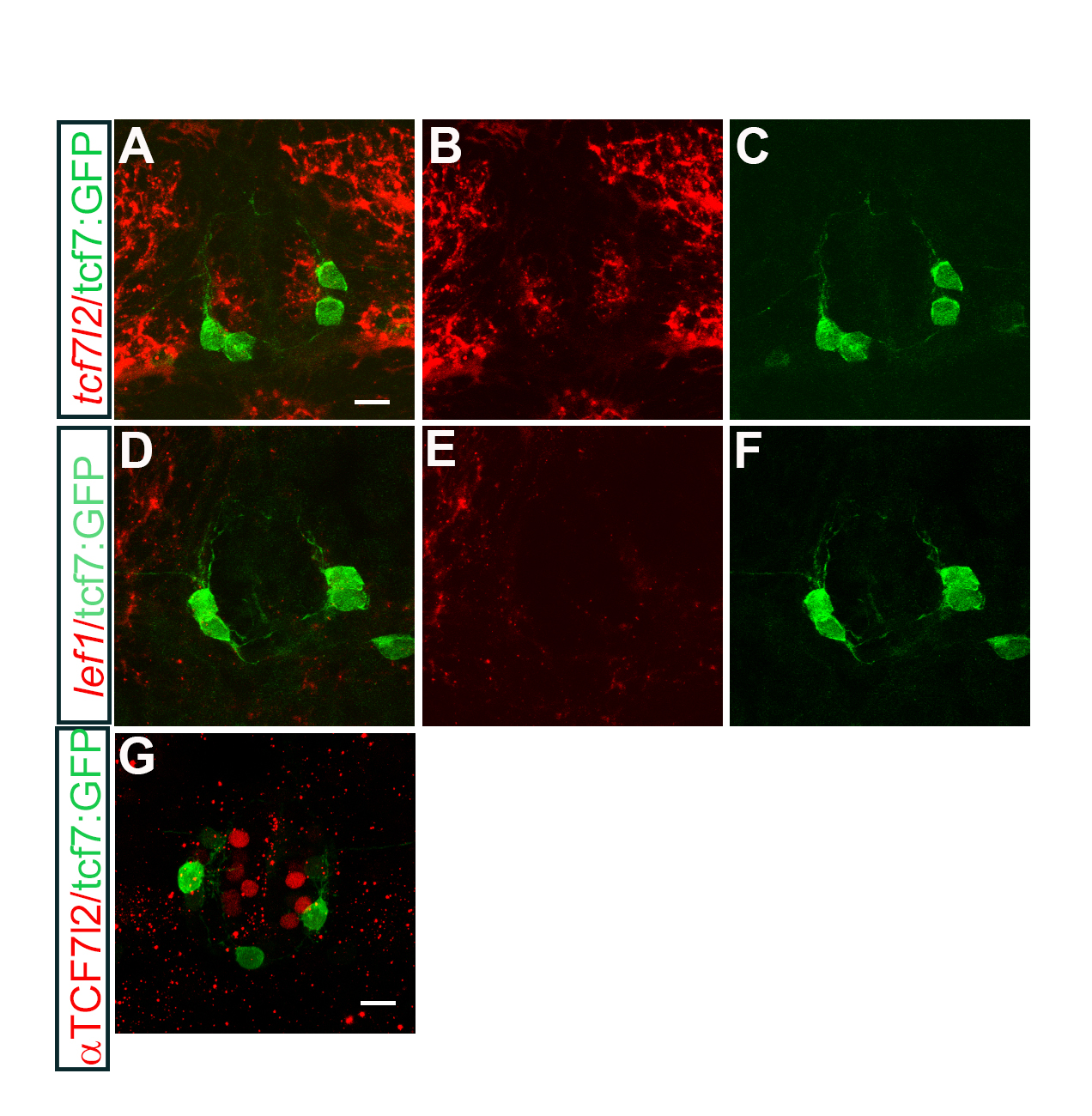

### Supplemental Figure 3

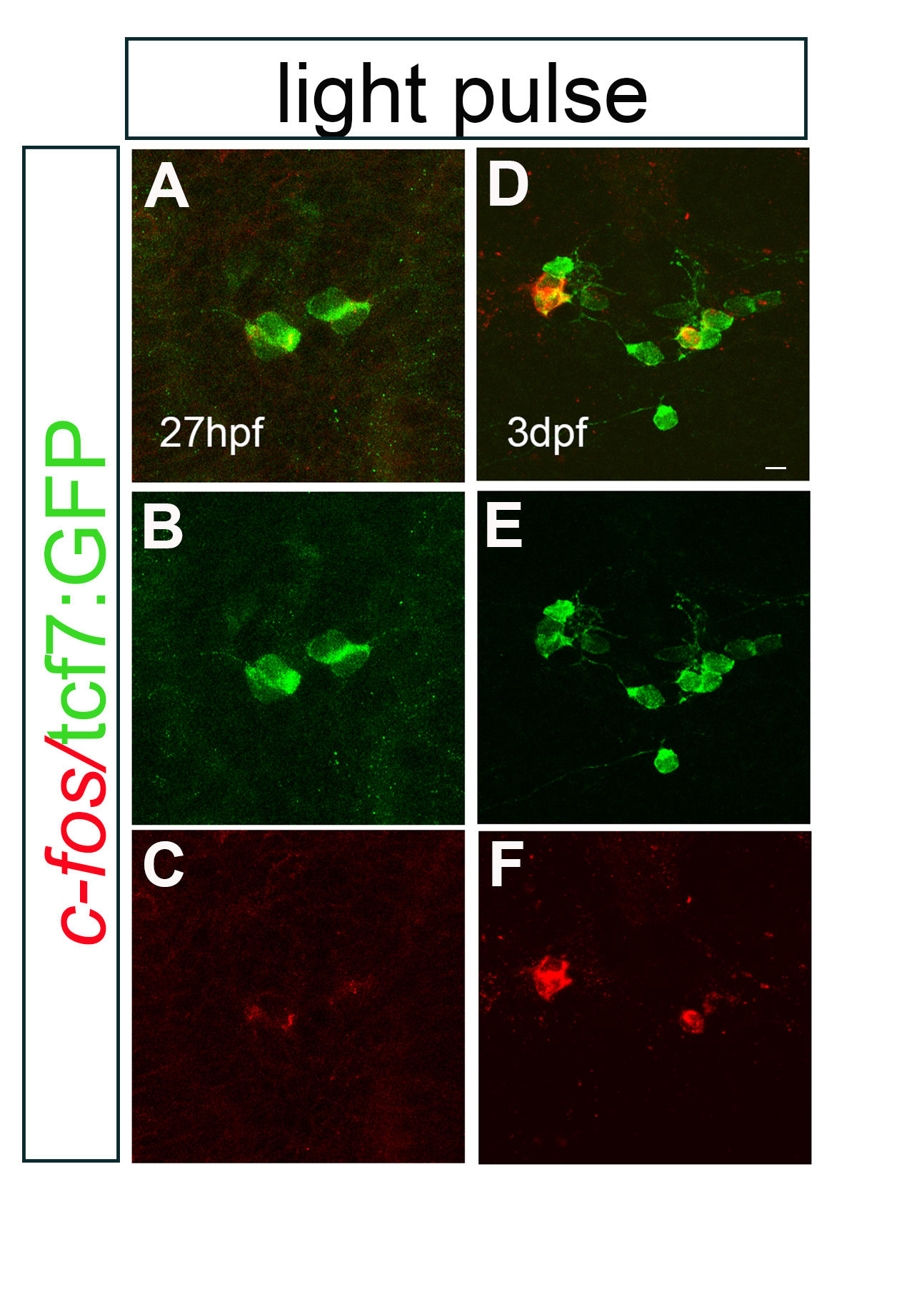
